# Dock5 is a new regulator of microtubule dynamic instability through GSK3β inhibition in osteoclasts

**DOI:** 10.1101/653055

**Authors:** Sarah Guimbal, David Guérit, Manon Chardon, Anne Blangy, Virginie Vives

**Affiliations:** Centre de Recherche de Biologie Cellulaire (CRBM), CNRS UMR 5237, 1919 route de Mende, 34293 Montpellier cedex 5, France; Montpellier University, 34095 Montpellier cedex 5, France

**Keywords:** Dock5, osteoclast, microtubule, knockout mice, resorption

## Abstract

**Background information:** Osteoclast resorption is dependent on a podosome-rich structure, called sealing zone, which is stabilized by acetylated microtubules. It tightly attaches the osteoclast to the bone creating a favorable acidic microenvironment for bone degradation. We already established that Rac activation by Dock5 is necessary for osteoclast resorption. Indeed, inhibition of Dock5 in osteoclasts results in Rac1 decreased activity associated to impaired podosome assembly into sealing zones and resorbing activity.

**Results:** In this report, we show that Dock5 knockout osteoclasts also present a reduced acetylated tubulin level leading to a decreased length and duration of microtubule growth phases whereas their growth speed remains unaffected. Dock5 does not act by direct interaction with the polymerized tubulin but through inhibition of the microtubules destabilizing kinase GSK3β downstream of Akt activation. Interestingly, we ruled out the implication of Rac1 in this process using specific inhibitors.

**Conclusion:** Our data involve Dock5 as a new regulator of microtubule dynamic instability in osteoclast.

**Significance:** The fact that Dock5 is a regulator of both actin cytoskeleton and microtubule dynamics makes it an interesting therapeutic target for osteolytic pathologies because of its dual role on sealing zone formation and stabilization.

## Introduction

Bone homeostasis is regulated through life by the balanced actions of highly specialized bone-forming osteoblasts and bone-resorbing osteoclasts. An important step of bone resorption is mineral phase dissolution by massive secretion of H^+^ and Cl^-^ ions. It exposes the collagen-rich underlying organic phase that becomes accessible to degradation by lysosomal proteases such as cathepsin K (Cappariello et al., 2014). To maintain this acidic microenvironment which is the key for an efficient resorption, osteoclasts tightly attach to the bone matrix by an actin-rich sealing zone composed of densely packed podosomes. Because of its nature, it was thought for a long time that its formation depended only on actin regulators whose roles have been extensively studied (Lowe et al., 1993) (Chellaiah et al., 2000) (Calle et al., 2004) (Destaing et al., 2008) (Croke et al., 2011) (Touaitahuata et al., 2014). A decade and a half ago, Jurdic’s lab discovery that an intact microtubule network was necessary for this adhesion structure stability (Destaing et al., 2003) emphasized the importance to understand the signaling pathways linking both cytoskeletons. On one hand, it was shown that belt stabilization is correlated to microtubule acetylation which is maintained by downregulation of two important players. First, RhoA-mDia2 dependent tubulin deacetylase HDAC6 activity (Destaing et al., 2005) is negatively controlled by Pyk2 (Gil-Henn et al., 2007). Independently, HDAC6 interaction with tubulin is antagonized by competitive binding of Cbl adaptor proteins (Purev et al., 2009). Second, the kinase GSK3β activity is inhibited by serine 9 phosphorylation by Akt. It allows stabilizing MAPs binding to growing microtubules (Matsumoto et al., 2013) and interaction between +TIP EB1 and actin belt associated cortactin (Biosse Duplan et al., 2014). On the other hand, it was shown that microtubules stabilize sealing zones through direct linkage mediated by actin regulators. Indeed, the unconventional myosin X encircles nascent sealing zones and provides force to expand them into mature expanded ones through interaction with microtubules (McMichael et al., 2010). More, the Dynamin 2 GTPase acts as an adaptor molecule between microtubules and actin podosomes, preventing them to scatter from the densely packed adhesion structure (Batsir et al., 2017). We showed that the Dock family member Dock5 is an exchange factor for the actin regulator Rac1 (Vives et al., 2011) (Ferrandez et al., 2017). During osteoclast resorption cycle, avβ3 integrins signaling induces Dock5 recruitment to the Src/Pyk2/p130Cas complex which ultimately leads to Rac1 activation and localization to the sealing zone (Vives et al., 2011) (Nagai et al., 2013). Dock5 plays an important role in this pathway because its deletion in knockout mice results in increased trabecular bone mass due to an impaired osteoclasts ability to assemble podosomes into sealing zones (Vives et al., 2011), a phenotype also observed in knockout mice for Rac1 (Croke et al., 2011) (Magalhaes et al., 2011) and its guanine exchange factors (Faccio et al., 2005) (Takegahara et al., 2010). Interestingly, a group working on allergy showed that Dock5 is essential for the dynamic rearrangement of microtubules during mast cell degranulation in anaphylactic responses (Ogawa et al., 2014).

In this study, we tested the hypothesis that Dock5 could play a central part in the control of osteoclast adhesion structure establishment through not only Rac1-dependent assembly (Vives et al., 2011) but also Rac1-independent formation/stabilization. We showed that Dock5 is indeed involved in sealing zone stabilization through microtubules dynamic instability regulation *via* the Akt-GSK3β pathway.

## Results

### Dock5^-/-^ osteoclasts present an abnormal microtubule acetylation pattern associated to a perturbed microtubule dynamic instability

Western blot analysis of proteins from wild-type and Dock5 knockout osteoclasts showed a significant decrease of tubulin acetylation in the absence of Dock5 (**figure 1A**). Confocal immunofluorescence microscopy allowed us to take a closer look at the microtubules distribution and acetylation pattern (**figure 1B**). As expected (Akisaka et al., 2011), microtubules of wild-type osteoclasts were typically organized (i) radially from the perinuclear region to the periphery where the podosome belt was located and (ii) circularly at the periphery. Both were modified with large evenly distributed acetylation patches. Although Dock5^-/-^ osteoclasts microtubules kept the radial orientation, the circular ones were missing at the periphery, which was consistent with the absence of podosome assembly into a belt. More, tubulin acetylation, that was present in small patches, was decreased in density compared to their wild-type counterpart (**figure 1C**).

**Figure 1:**
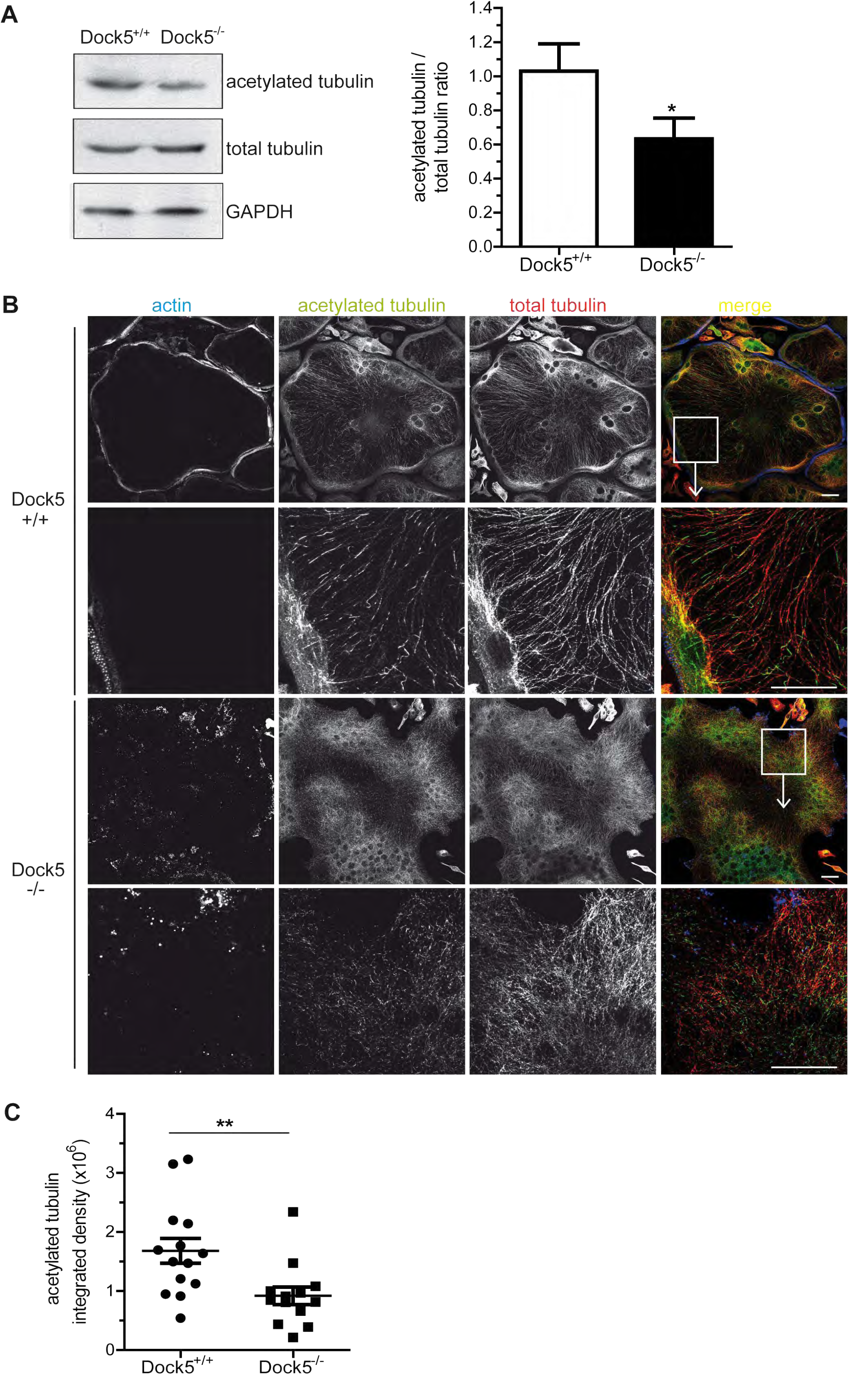
Microtubule acetylation in Dock5^-/-^ osteoclasts. **(A)** Representative western blot showing acetylated tubulin, total tubulin and GAPDH in Dock5^+/+^ and Dock5^-/-^ osteoclasts (left). Acetylated tubulin levels from 7 independent experiments were normalized to total tubulin and represented on a bar graph (right). *, p<0.05 using one-tailed Mann Whitney test. **(B)** Actin (blue), acetylated (green) and total (red) tubulin immunofluorescence staining of Dock5^+/+^ and Dock5^-/-^ osteoclasts. For each genotype, bottom panels show an enlarged view of the top panels boxed areas. Scale bar= 20 μm. **(C)** Graph showing acetylated tubulin integrated density from immunofluorescent staining of 13-14 Dock5^+/+^ and Dock5^-/-^ osteoclasts from different experiments. **, p<0.01 as determined by twotailed Mann Whitney test.

Since tubulin acetylation is associated with microtubule stabilization (Matsumoto et al., 2013), we reasoned that this acetylation defect could affect their dynamic instability. To answer this question, we expressed the EB3 +TIP fused to GFP in wild-type and Dock5 knockout osteoclasts to visualize their growing microtubule plus ends using time-lapse video microscopy (**supplementary figure 1**). Tracking of EB3-GFP at the +TIP of microtubules (**figure 2A**) showed a significant decrease of microtubule polymerization duration (**figure 2B**) and length (**figure 2C**) in the absence of Dock5. Polymerization speed however remained unchanged (**figure 2D**). More, the mean total number of actively growing microtubules was similar for both genotypes per image of the movies (**figure 2E**) but the total comet tracks number was significantly increased in Dock5^-/-^ osteoclasts over a 1 min period (**figure 2F**), suggesting an increased number of catastrophes.

**Figure 2:**
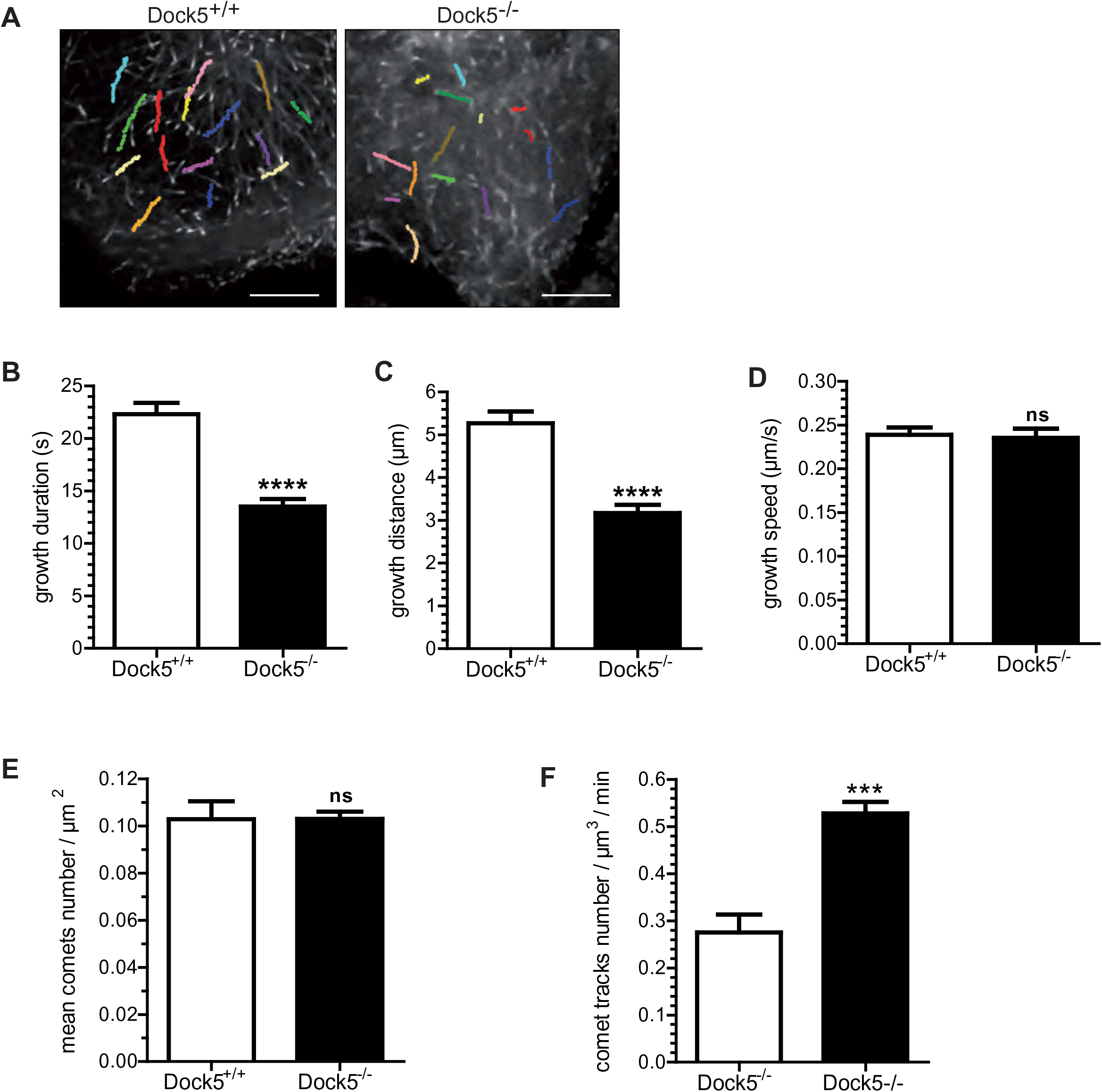
Microtubule dynamic instability in Dock5^-/-^ osteoclasts. **(B)** GFP-positive comets were tracked in Dock5^+/+^ and Dock5^-/-^ osteoclasts infected with EB3-GFP. Scale bar= 20 μm. Bar graphs showing microtubule growth **(B)** duration, **(C)** distance and **(D)** speed as well as **(E)** comets number / μm^2^ and **(F)** comet tracks number / μm^3^ / min of 18-19 osteoclasts from 4 independent experiments. **(B-F)** ns, not significant; *, p<0.05; ***p<0.0001; ****p<0.00001 as determined by two-tailed Mann Whitney test.

### Dock5 regulates microtubules dynamics through a Rac-independent pathway

It has been shown that Dock5 is a guanine exchange factor for Rac1 whose activity is decreased in Dock5 knockout osteoclasts (Vives et al., 2011). That made us wonder if the microtubule phenotype we described in Dock5^-/-^ osteoclasts was Rac-dependent. We reasoned that, if it is the case, we should reproduce it when we specifically inhibit Dock5-dependent Rac activation in wild-type osteoclasts. Western blot analysis showed that, opposite to what was observed in Dock5 knockouts, treatment with an allosteric inhibitor of Dock5 called C21 (Vives et al., 2011) (Vives et al., 2015) (Ferrandez et al., 2017) (Xu et al., 2017a) significantly increased tubulin acetylation level (**figure 3A**). It was confirmed by confocal immunofluorescence microscopy that showed that, in addition to the podosome belt disruption, C21 treatment induced a more intense microtubules acetylation compared to wild-type osteoclasts (**figure 3B**). As expected, we observed the same phenotype using EHT 1864, a compound that prevents Rac guanine nucleotide association and consequently inhibits downstream signaling (Shutes et al., 2007). Indeed, tubulin acetylation augments with increasing concentrations of the drug in osteoclasts (**figure 4A**) that show a disrupted microtubule network (**figure 4B**). In order to check if the acetylation alteration affected microtubules dynamics, we treated or not EB3-GFP expressing wild-type osteoclasts with C21 and recorded the resulting comets using time-lapse video microscopy. Their tracking showed a significant decrease of microtubule polymerization duration, length and speed after C21 treatment (**figure 3C**). More, C21 impacted neither the total number of actively growing microtubules on each image of the movies (**supplementary figure 2A**) nor the comet tracks number over a 1 min period (**supplementary figure 2B**). These results show that actin belt disruption following the complete (EHT 1864) or partial (C21) inhibition of Rac activity does not reproduce the microtubule phenotype observed in Dock5 knockouts. It suggests that Dock5 regulates microtubules independently of its Rac guanine exchange factor activity in osteoclast.

**Figure 3:**
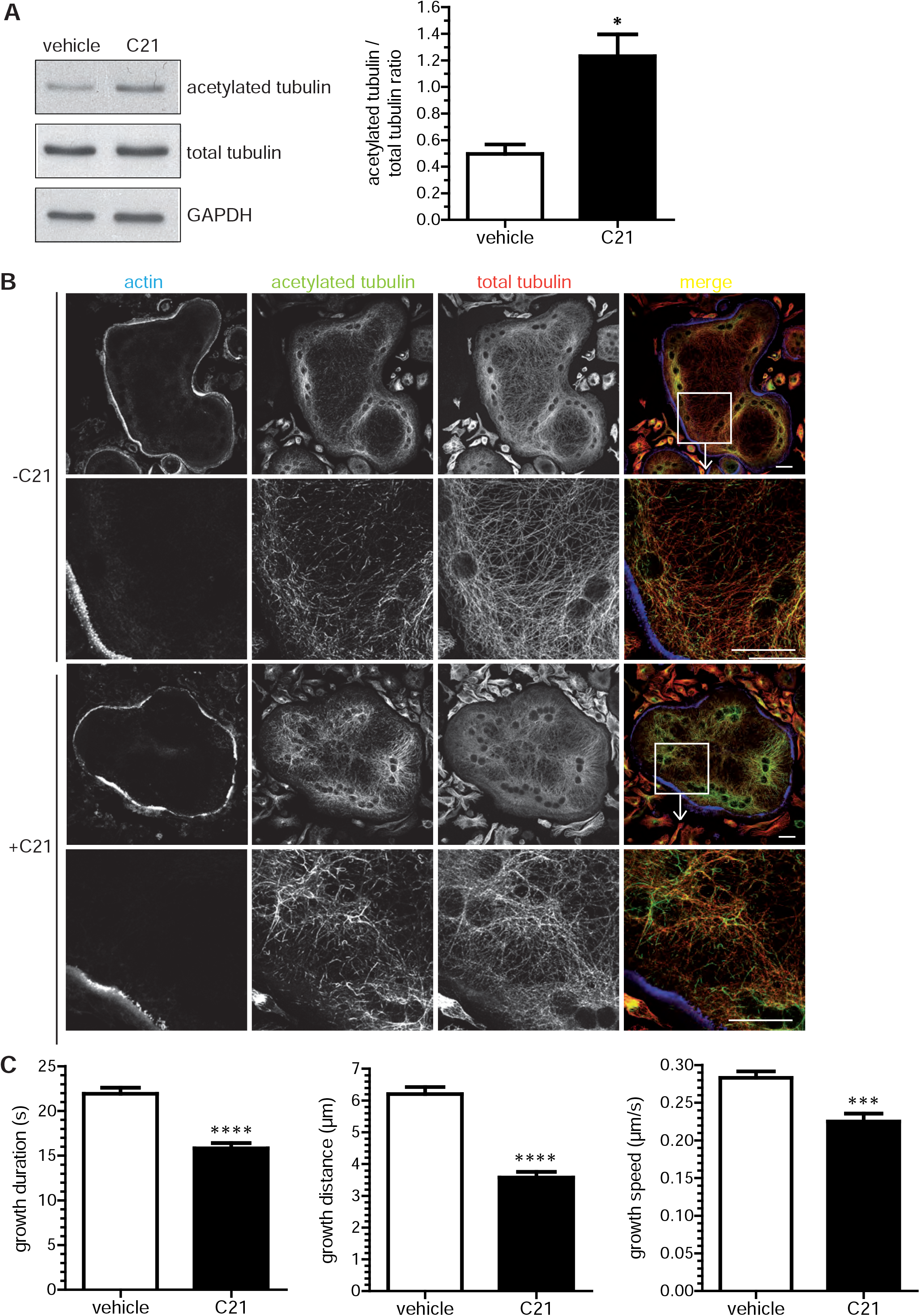
C21 effect on microtubule acetylation and dynamic instability. **(A)** Representative western blot showing acetylated tubulin, total tubulin and GAPDH in wild-type osteoclasts treated (C21) or not (vehicle) with 100 μM C21 for 1 hour (left). Acetylated tubulin levels from 6 independent experiments were normalized to total tubulin and represented on a bar graph (right). **(B)** Actin (blue), acetylated (green) and total (red) tubulin immunofluorescence staining of Dock5^+/+^ osteoclasts treated (+C21) or not (-C21) with 100 μM C21 for 1 hour. For each condition, bottom panels show an enlarged view of the top panels boxed areas. Scale bar= 20 μm. **(C)** Bar graphs showing microtubule growth duration, distance and speed in Dock5^+/+^ osteoclasts expressing EB3-GFP and treated (C21) or not (vehicle) in the same conditions as above in 14-20 osteoclasts from 3 independent experiments. (A,C) *, p<0.05; ***, p<0.0001; ****, p<0.00001 as determined by two-tailed Mann-Whitney test.

**Figure 4:**
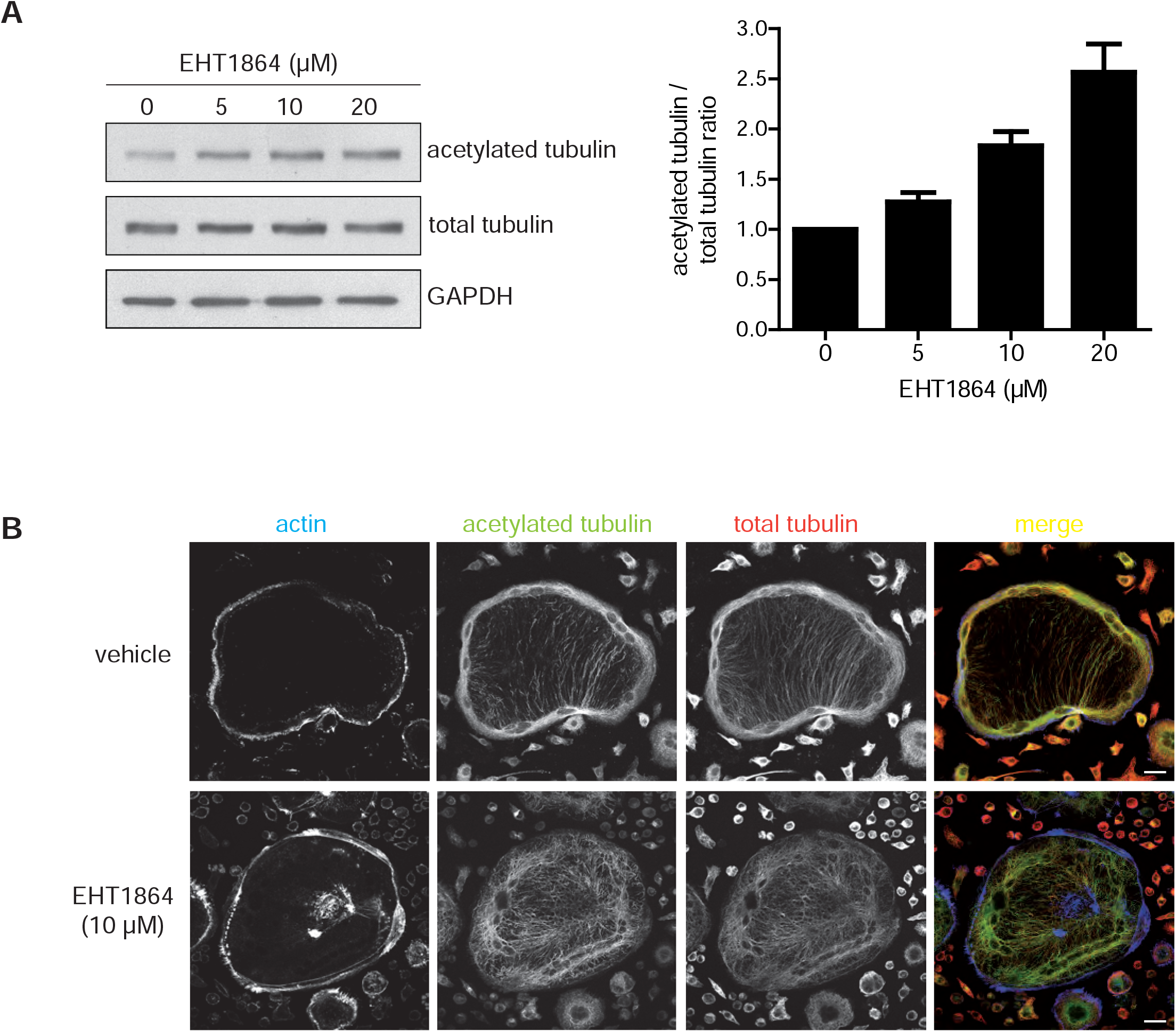
Rac inhibitor EHT1864 effect on microtubule acetylation. **(A)** Representative western blot showing acetylated tubulin, total tubulin and GAPDH in wild-type osteoclasts treated with increasing concentrations of EHT1864 for 2 hours (left). Acetylated tubulin mean level from 3 independent experiments ±sem were normalized to total tubulin and represented on a bar graph (right). **(B)** Actin (blue), acetylated (green) and total (red) tubulin immunofluorescence staining of Dock5^+/+^ osteoclasts treated (EHT1864) or not (vehicle) with 10 μM EHT1864 for 2 hours. Scale bar= 20 μm.

### Dock5 regulates microtubules through a distant signaling pathway

Consistently with its Rac-activating role, we previously showed that Dock5 colocalizes with vinculin, an actin cloud-associated protein, at the podosome belt in RAW264.7 cell-derived osteoclasts (Vives et al., 2011). Since we uncovered a new microtubule-regulating function, we wondered whether a Dock5 protein fraction could also be associated to microtubules. To answer this question, we separated soluble and polymerized microtubules. Western blot analysis showed that Dock5 was not associated to the polymerized fraction, which contained acetylated tubulin and could be depolymerized upon nocodazole treatment (**figure 5A**). These results suggest that Dock5 microtubules regulation is not mediated by direct association but through a distant signaling pathway.

**Figure 5:**
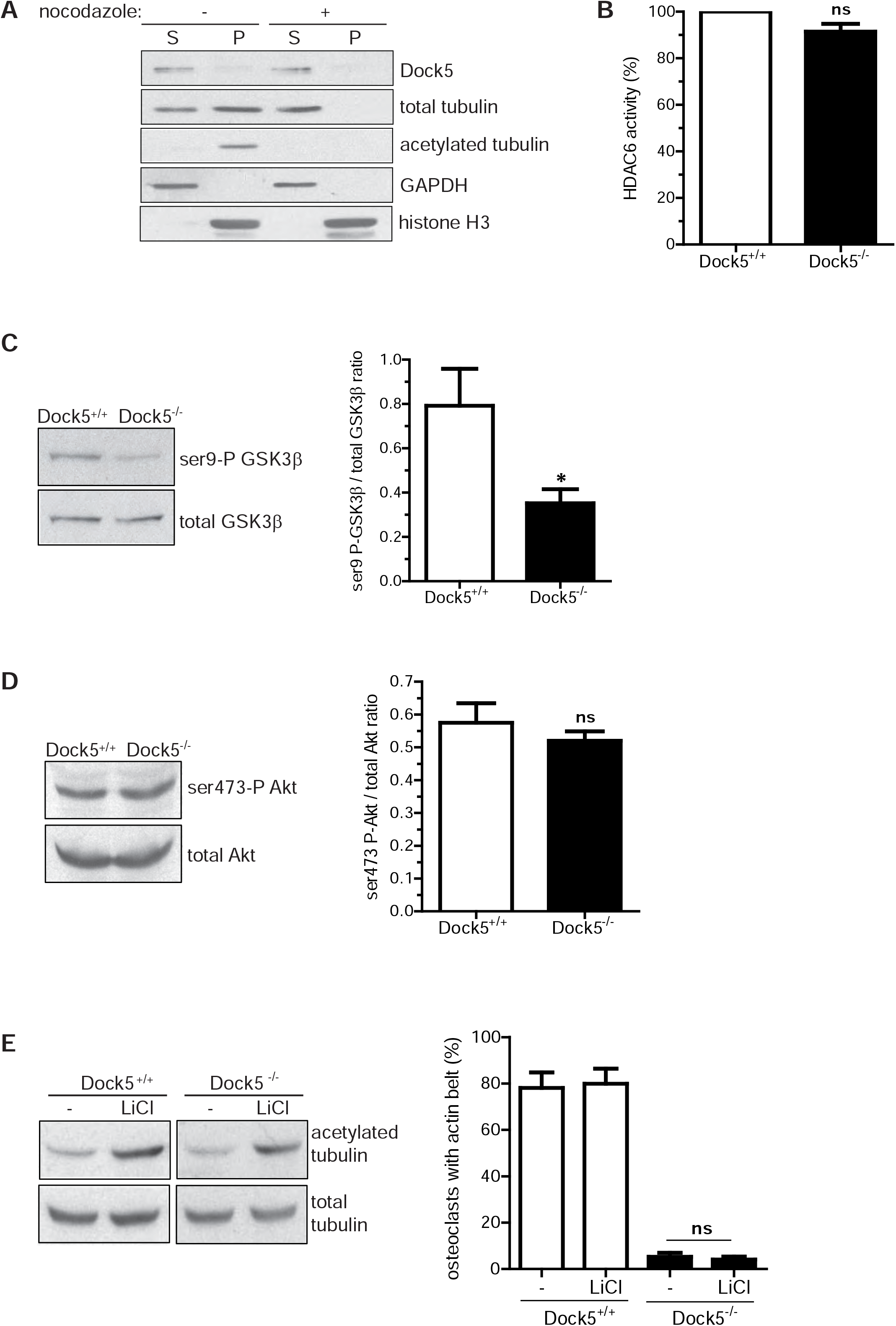
Defective signaling pathways in Dock5^-/-^ osteoclasts. **(A)** Western blot showing Dock5 association to soluble (S) or polymerized (P) tubulin from wild-type osteoclasts treated (+) or not (-) with 10 μM nocodazole for 1 hour. **(B)** HDAC6 activity in Dock5^+/+^ and Dock5^-/-^ osteoclasts. ns, not significant as determined by Wilcoxon Signed Rank test. Representative western blots showing **(C)** GSK3β- and **(D)** Akt phosphorylation levels in Dock5^+/+^ and Dock5^-/-^ osteoclasts (left). Signals from 3-5 independent experiments were normalized to the total protein levels (right). ns, not significant; *, p<0.05 as determined by two-tailed Mann-Whitney test. **(E)** Dock5^+/+^ and Dock5^-/-^ osteoclasts were treated (LiCl) or not (-) with 20 mM lithium chloride for 5 hours and tubulin acetylation was observed by western blot (left). The treatment effect on actin belt formation/stabilization was quantified from 3 independent experiments (right). ns, not significant as determined by two-tailed Mann Whitney test.

### Dock5^-/-^ osteoclasts show an increased GSK3β activity

In order to identify the mechanism by which the microtubules were destabilized in Dock5 knockouts, we searched for deregulations of pathways involved in microtubules acetylation in osteoclasts. HDAC6 is a deacetylase whose activity is kept to a minimum (Gil-Henn et al., 2007) (Purev et al., 2009) to allow a stable microtubule network. We therefore checked whether HDAC6 activity was higher Dock5^-/-^ osteoclasts but we could not find any differences between genotypes (**figure 5B**). We next focused on the Akt-GSK3β axis. Akt inhibits GSK3β; a serine threonine kinase GSK3β which destabilizes microtubules by regulating microtubules associated proteins interactions (Matsumoto et al., 2013). Western blot analysis of GSK3β inhibitory serine 9 phosphorylation showed that GSK3β activity was increased in Dock5^-/-^ osteoclasts (**figure 5C**). Interestingly, Akt activity was not modified (**figure 5D**), suggesting that Dock5 acts downstream in this pathway.

Mature Dock5^-/-^ osteoclasts mainly present isolated or clustered podosomes as if they were not able to assemble or maintain them into a belt (Vives et al., 2011). Since acetylated microtubules were shown to be necessary for podosome belt assembly/stabilization (Destaing et al., 2003) (Destaing et al., 2005), we assumed that restoration of a normal acetylation level in Dock5^-/-^ osteoclasts would rescue the actin belt. Therefore, wild-type and Dock5 knockout osteoclasts were cultured in the presence of GSK3β inhibitor lithium chloride. Although the treatment increased tubulin acetylation at a similar level between the two genotypes, it was not enough to allow the actin belt formation/stabilization in the absence of Dock5 (**figure 5E**). The same result was obtained using CHIR99021, a more selective GSK3β inhibitor **(data not shown)**.

There is a close relationship between microtubules and the actin cytoskeleton in osteoclasts. Microtubules grow toward isolated/clustered podosomes during osteoclast differentiation and then form a ring at the cell periphery where the podosome belt is located (Akisaka et al., 2011). During this process, they are necessary for podosomes transition from rings to belt (Destaing et al., 2003). To our knowledge, no data have been made available about the microtubule network in mice knockout for actin regulators, but we would expect an effect on the microtubules dynamics. Since the actin regulator Rac activity is also decreased in Dock5^-/-^ osteoclasts (Vives et al., 2011), we decided to reestablish not only GSK3β inhibition but also Rac activation hypothesizing that both might be needed for a wild-type podosome belt formation/stabilization. Contrary to our expectations, lithium chloride treatment of Rac L61 infected knockout osteoclasts did not rescue the phenotype **(data not shown)**. However, we cannot exclude the fact that a better controlled regulation of Rac activation level and/or localization is important for this process.

## Discussion

Osteoclast resorption function depends on podosome assembly into sealing zones. More and more evidence show now that the interplay between the actin cytoskeleton and microtubules is crucial for this adhesion structure formation/stabilization but the overall mechanism is still poorly understood. We already established that Dock5 is essential for sealing zones formation through Rac-dependent actin regulation. In this report, we show that Dock5 might also be involved in its stabilization by controlling microtubule dynamic instability in a Rac-independent pathway. Dock5 deletion results in decreased microtubules growth length and duration associated to reduced acetylated tubulin level. This post translational modification is a hallmark of stability because it increases mechanical resilience to ensure the persistence of long-lived microtubules (Xu et al., 2017b) (Janke and Montagnac, 2017). Its downregulation, resulting from either HDACs inhibition or GSK3β activation, has been correlated to microtubule dynamics alteration in different cell types (Garza et al., 2018) (Bacon et al., 2015) (Qu et al., 2017).

Dock5 belongs to the Dock family of guanine nucleotide exchange factors which are atypical activators of small G proteins through evolutionarily conserved DHR2 domain (Laurin and Côté, 2014). Accordingly, Dock5 published functions are mainly related to Rac activation (Omi et al., 2008) (Vives et al., 2015) (Vives et al., 2011) (Ferrandez et al., 2017) (Biswas et al., 2019) in various cellular processes such as neutrophil chemotaxis and superoxide production(Watanabe et al., 2014), epithelial cell invasion and metastasis (Frank et al., 2017) or lens protection (Xu et al., 2017a) (Omi et al., 2008). Ogawa *et al.* were the first ones to describe a Rac-independent role in microtubule network remodeling during mast cell degranulation. In these cells, Dock5 acts as a key signaling adaptor that brings Akt into the vicinity of GSK3β, allowing it to phosphorylate GSK3β on serine 9 and inhibit its activity (Ogawa et al., 2014). The fact that Akt activation remains unchanged whereas that of GSK3β is increased in Dock5 knockout osteoclasts suggests that the same mechanism takes place in both cell types. Ubiquitously-expressed serine threonine kinase GSK3β is a critical negative regulator of microtubule dynamic instability (Xu et al., 2015) (Kumar et al., 2009) (Barnat et al., 2016). It phosphorylates microtubule-binding proteins such as MAP1B (Barnat et al., 2016), CRMP-2 (Liz et al., 2014) (Garza et al., 2018), CLASP-2 (Kumar et al., 2009) (Schmidt et al., 2012), Tau (Xu et al., 2015) (Gąssowska et al., 2014) or APC (Asada and Sanada, 2010) (Zhou et al., 2004), preventing them from stabilizing microtubules. Some of them are recruited by EB1 at growing microtubule ends that can be assimilated to a molecular platform establishing the link between microtubules and other structures (Akhmanova and Steinmetz, 2015). In osteoclasts, EB1 is enriched in microtubules that are directed toward podosomes (Akisaka et al., 2011). Depletion of Akt results in EB1/APC loss of interaction leading to reduced sealing zone and bone resorption, which are recovered upon GSK3β inhibition (Matsumoto et al., 2013). More, EB1 which is indispensable for podosome patterning, interacts with podosome-containing acetylated cortactin in the belt (Biosse Duplan et al., 2014). It would be interesting to test if either one of these proteins phosphorylation status or localization is impacted in the absence of Dock5.

Our observations strongly suggest that Dock5 is necessary for osteoclast adhesion structure establishment by stimulating dual pathways: Rac1-dependent podosome formation/patterning and Rac1-independent stabilization *via* Akt-dependent GSK3β inhibition (**figure 6**). In the absence of Dock5, decreased Rac activation and GSK3β inhibition are observed. Since both are involved in podosome belt formation and stabilization respectively, we were surprised that a simultaneous recovery of their physiological levels could not rescue podosome belts in Dock5^-/-^ osteoclasts. This process might need a finely tuned amount of Rac activation and/or localization that can hardly be achieved though primary cells infection. Similarly, although lithium chloride treatment increases *in fine* microtubule acetylation level, the latter is much higher than in control osteoclasts (**figure 5E**). That might affect the expected rescue because both positive and negative regulation of acetylation impact microtubules growth characteristics (Qu et al., 2017) (Zilberman et al., 2009). Nevertheless, further data are consistent with our model. Among them, the fact that partial (C21) or complete (HT 1864) inhibition of Rac1 activity did not reproduce Dock5^-/-^ osteoclasts phenotype suggesting that at least part of it is independent from this GTPase. In the same line of idea, since Rac1 signaling pathway has been shown to downregulate Op18 catastrophe promoting activity through its phosphorylation (Watabe-Uchida et al., 2006) (Wittmann et al., 2004), we wondered whether Op18 phosphorylation levels were decreased in Dock5^-/-^ osteoclasts which show a reduced Rac activity (Vives et al., 2011). Interestingly, they were not modified by the absence of Dock5 expression in osteoclasts (unpublished data). Dock5 is not the only member of the Dock family to regulate microtubules independently from its guanine exchange factor activity. Dock7 controls the genesis of neurons from radial glial progenitor cells during cortical development through antagonistic interaction with the microtubule growth-promoting function of TACC3 (Yang et al., 2012). And finally, Dock3 participates in BDNF-dependent primary hippocampal neurons axonal outgrowth by also regulating (i) Rac1-dependent actin reorganization and (ii) microtubule assembly through GSK3 binding, recruitment to the plasma membrane and inactivation. The resulting decrease in CRMP-2 and APC phosphorylation allows their association to tubulin dimers and promotes microtubules polymerization (Namekata et al., 2012).

**Figure 6:**
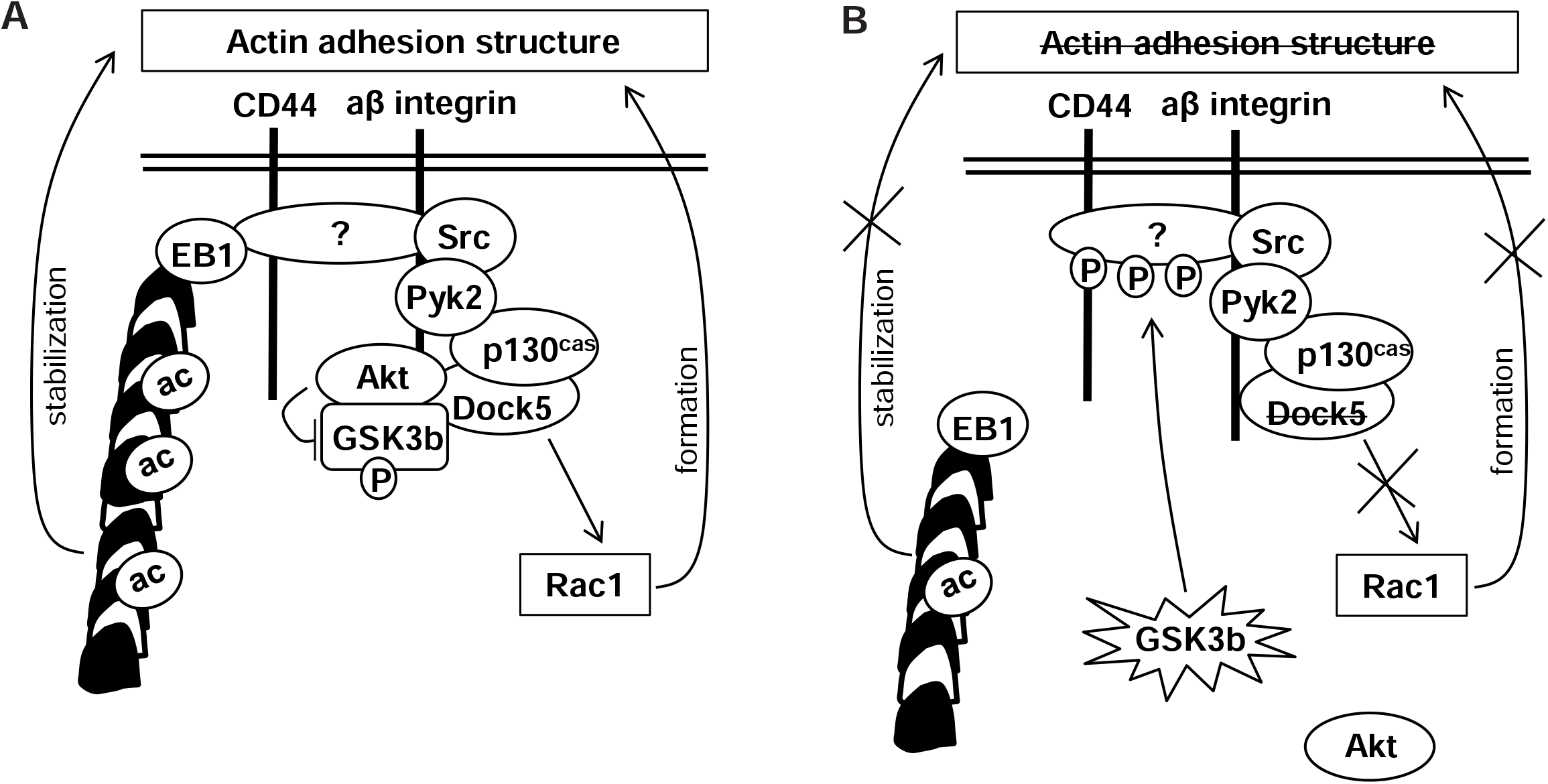
Proposed role of Dock5 in osteoclast adhesion structure establishment. **(A)** αβ integrin activation induces Doc5 recruitment to the Src/Pyk2/p130Cas complex leading to Rac1 activation and podosome assembly and patterning. In addition, Dock5 binding to Akt brings the latter sufficiently close to GSK3β to inactivate it by phosphorylation on serine 9. EB1 can interact with unidentified (?) protein(s) associated to podosomes allowing the acetylated microtubules to stabilize the actin structure. **(B)** In the absence of Dock5, decreased Rac1 activation prevents efficient podosome formation. More, activated GSK3β phosphorylates unidentified (?) protein(s) hindering its / their interaction with EB1-decorated microtubules that are destabilized.

Altogether, our data involve Dock5 as a new regulator of microtubule dynamic instability in osteoclast making it is an interesting therapeutic target for osteolytic pathologies because of its dual role on sealing zone formation/patterning and stabilization.

## Materials and Methods

### Ethics statement

Mice sacrifice and bone marrow harvest were performed in compliance with local animal welfare laws, guidelines and policies, according to the rules of the regional ethical committee.

### Reagents

C21 (N-(3,5-dichlorophenyl)-benzenesulfonamide) was synthetized by Roowin, lithium chloride was purchased from Fluka (#62476), EHT1864 from Santa Cruz (#sc-361175) and nocodazole from Sigma (#M1404).

Bone Marrow Macrophages (BMMs) isolation and differentiation in osteoclasts Bone marrow cells were purified from long bones of 6- to 8- week-old C57BL/6 Dock5 wild-type and knockout mice sacrificed by cervical dislocation. 3.10^6^ bone marrow cells were cultured for 3 days in αMEM containing 10% heat-inactivated fetal calf serum (Biowest), 2 mM glutamine and 100U/ml penicillin-streptomycin supplemented with 100 ng/ml M-CSF (Miltenyi, #130-101-703) in 100mm-diameter petri dishes. Medium was changed and the cells cultured for another 3 days to obtain BMMs. For osteoclast differentiation, 3.10^4^ BMMs/well were seeded in 24-well tissue culture plates and cultured with 30 ng/ml M-CSF and 50 ng/ml RANKL (Miltenyi, #130-094-076) for 4-6 days. Media was changed every 2 days.

### Retroviral infection of BMMs

pMXS-EB3-GFP or pMXS-Rac L61-GFP was co-transfected with pC57GP-Gag-pol and pCSIG-VSV-G constructs into Hek 293T cells using Jet-PEI (Polyplus transfection, #101-10 N). Six hours later, the medium was replaced with fresh one and cells were grown for an additional 72 h. The conditioned medium containing recombinant retroviruses was collected and filtered through 0.45 μm-pore-size filters. BMMs were cultured with fresh retroviral supernatant diluted to half in αMEM containing 10% fetal calf serum, 2 mM glutamine, 8 μg/ml polybrene (Sigma) and 100 ng/ml M-CSF for 24 h and then selected for expression of the construct with 3 μg/ml puromycin (Sigma) for 24 h. Puromycin-resistant BMMs were used for osteoclast differentiation.

### Time-lapse microscopy and microtubules dynamic instability analysis

EB3-GFP expressing osteoclasts were placed on a 37°C heated stage in a 5% CO_2_ chamber. Images were acquired using a Yokogawa CSU-X1 spinning-disk confocal unit (Andor Technology) and an EMCCD iXon Ultra camera coupled to a Nikon Ti Eclipse microscope (Nikon Instruments) controlled by the Andor iQ3 software (Andor Technology). They were collected every second after 488-nm laser illumination using a 60X plan apo lambda 1.4 NA objective over a 2 min period and processed using ImageJ PureDenoise plugin. EB3-GFP comets appearing in at least 3 consecutive sections were manually tracked with ImageJ MtrackJ plugin. Comet count and comet tracks count were obtained using an ImageJ custom-written macro.

### HDAC6 activity measurement

HDAC6 activity was assessed using the Fluorogenic HDAC6 assay kit (Bioscience, #50076) according to the manufacturer’s instructions.

### Soluble and polymerized tubulin fractionation

Soluble and polymerized tubulin was fractionated as described (Larrieu et al., 2014). Briefly, osteoclasts pre-treated or not with 10 μM nocodazole for 2 h were lysed in Microtubules Stabilization Buffer (MSB: 85 mM PIPES pH 6.9, 1 mM EGTA, 1 mM MgCl_2_, 2M glycerol, 0.5% triton, 4 μg/ml taxol) supplemented with proteases inhibitors. Lysates were kept in ice for 2 min and then centrifuged for 10 min at 13000 rpm. Laemmli buffer was added to the supernatants (soluble fraction). Pellets (polymerized fraction of proteins) were washed once in MSB without triton and resuspended in Laemmli buffer.

### Western Blot

Whole osteoclast extracts were made in Laemmli buffer, resolved on SDS-PAGE and electrotransferred on nitrocellulose membranes. Immunoblotting was performed using the following primary antibodies: mouse anti-lys40 acetylated alpha tubulin (1/1000, Sigma, #T6793), mouse anti-alpha tubulin (1/1000, Sigma, #T6074), rabbit anti-GAPDH (1/2000, Cell Signaling, #2118), rabbit anti-EB1 (1/2500, Sigma, #E3406), rabbit anti-ser9 phospho GSK3β (1/1000, Cell Signaling, #9322), rabbit anti-GSK3β (1/1000, Cell Signaling, #12456), rabbit anti-ser473 phospho Akt (1/1000, Cell Signaling, #4060), rabbit anti-Akt (1/1000, Cell Signaling, #9272), rabbit anti-ser16 phospho Op18 (1/500, Santa Cruz, #sc-12948-R), rabbit anti-Dock5 (1/2000, homemade) and rabbit anti-histone H3 (1/5000, Abcam, #1791). Signals were visualized by the ECL Western Lightning Plus detection system (Perkin Elmer) with horseradish peroxidase-conjugated secondary antibodies (GE Healthcare) and quantified using ImageJ.

### Immunofluorescence and microscopy

Osteoclasts differentiated on 13 mm-diameter glass coverslips were fixed for 20 min, in 10 μM Taxol (Sigma, #T7402) and 3,2% paraformaldehyde in PHEM (60 mM Pipes, 25 mM Hepes, 10 mM EGTA, 4 mM MgSO_4_, pH 6.9). After permeabilization with 0.1% Triton X100 for 1 min and blocking with 1% BSA in PBS for 15 min, osteoclasts were incubated for 1 h with Alexa Fluor 647-labelled phalloidin (1/1000, Life Technologies, #A22287), mouse anti-lys40 acetylated alpha tubulin (1/500, Sigma, #T6793), rat anti-alpha tubulin (1/200, Santa Cruz, #sc-53030) or rabbit anti-Dock5 (1/100, homemade) and revealed with the adapted Alexa Fluor 488 or 546-conjugated secondary antibodies (1/1000, Life Technologies). Preparations were mounted in Citifluor mouting medium (Biovalley) and imaged with Leica SP5-SMD confocal microscope using 40X HCX Plan Apo CS oil 1.3NA or 63X HCX Plan Apo CS oil 1.4NA objectives. All imaging was performed at the Montpellier RIO Imaging facility (http://www.mri.cnrs.fr/en/). Immunofluorescent signals integrated density was quantified using ImageJ.

### Statistical analysis

Graphs are represented as the mean ± sem. Statistical significance was assessed with the non-parametric statistical tests mentioned in figure legends using GraphPad Prism (GraphPad Software, Inc).

---

**Supplementary figure 1: EB3-GFP comets in Dock5^+/+^ and Dock5^-/-^ osteoclasts** Representatives movies with 1 frame per second showing EB3-GFP comets from **(A)** Dock5^+/+^ and **(B)** Dock5^-/-^ osteoclasts.

**Supplementary figure 2: EB3-GFP comets analysis in C21-treated wild-type osteoclasts** Bar graphs showing (A) comets number / μm^2^ and (B) comet tracks number / μm^3^ / min in wild-type osteoclasts treated (C21) or not (vehicle) with 100 μM C21 for 1 hour in 3 independent experiments. ns, not significant as determined by two-tailed Mann-Whitney test.

## Acknowledgments

We acknowledge the Montpellier RIO Imaging (https://www.mri.cnrs.fr/en/) and the RAM (http://www.ram.cnrs.fr/) facilities. We thank J. Boudeau (CRBM, Montpellier, France) for EB3-GFP expressing vector, Carlos Sanchez-Huertas (CRBM, Montpellier, France) for helpful discussions and Anne Morel (CRBM, Montpellier, France) for the management of Dock5 transgenic mice colonies.

## Abbreviation Meaning

APC: Adenomatous polyposis coli
BMM: Bone Marrow Macrophages
GSK3β: Glycogen Synthase Kinase 3 beta
HDAC6: Histone Deacetylase 6

