## Supplementary figures and images for "Dock5 is a new regulator of microtubule dynamic instability through GSK3β inhibition in osteoclasts"

### Supplemental figure 2

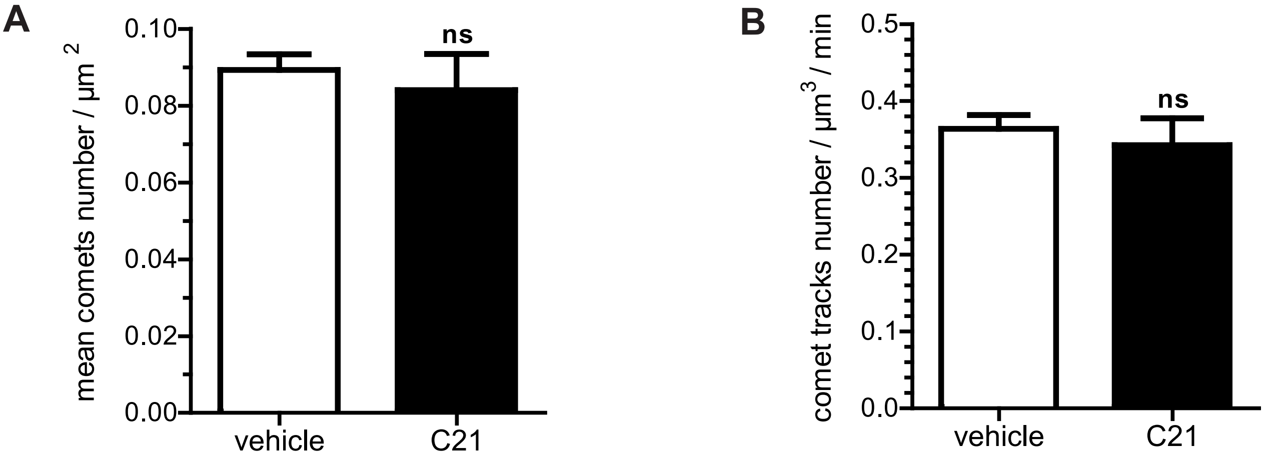


Supplementary figure 2
